# N-terminal acetylation reduces α-synuclein pathology in models of Parkinson’s disease

**DOI:** 10.1101/2025.11.21.689703

**Authors:** Esther del Cid-Pellitero, Thomas Goiran, Zaid A.M. Al-Azzawi, Nathan C. Karpilovsky, Jace Jones-Tabah, Jonas J. Mayo, Frederique Larroquette, Wen Luo, Irina Shlaifer, Yogitha Thattikota, Andrew Bayne, Jean-Francois Trempe, Thomas M. Durcan, Edward A. Fon

## Abstract

The α-synuclein protein, encoded by *SNCA* gene, is a major constituent of pathological intracellular inclusions such as Lewy bodies found in the brains of patients with Parkinson’s disease and other synucleinopathies. Whereas α-synuclein phosphorylation has been much studied, comparatively less work has been devoted to other post-translational modifications such as acetylation, especially given that N-terminally acetylated α-synuclein is the most abundant endogenous form of the protein in the brain. In this study, using multiple *in vitro* and *in vivo* models, we sought to better understand the role of N-terminal acetylation in the pathogenesis of synucleinopathies. We found that N-terminal acetylation slowed aggregation of both α-synuclein monomers and pre-formed fibrils *in vitro*. Uptake of acetylated α-synuclein pre-formed fibrils into both immortalized cell lines and iPSC-derived dopamine neurons was also slowed compared non-acetylated fibrils. In addition, exposure to acetylated pre-formed fibrils induced less seeding of endogenous α-synuclein, as measured by the accumulation of Serine129-phosphorylated α-synuclein inclusions in both iPSC-derived dopamine neurons and mouse brain. Finally, mice injected with N-terminally acetylated α-synuclein pre-formed fibrils survived significantly longer than mice injected with non-acetylated fibrils. Taken together, our study indicates that N-terminal acetylation reduces α-synuclein aggregation, uptake into cells, seeding of endogenous α-synuclein, and toxicity *in vivo*, suggesting that this prevalent post-translational modification represents a potent, physiologically relevant protective mechanism, which has thus far largely not been taken into consideration in most experimental paradigms of Parkinson’s disease and synucleinopathies.

## Introduction

One of the main pathological hallmarks of Parkinson’s disease (PD) and other synucleinopathies is the presence of intracellular inclusions such as Lewy bodies, consisting of aggregated and phosphorylated α-synuclein (α-syn) protein along with accumulated lipids and organelles ^1–3^. Moreover, mutations and copy number variants in the *SNCA* gene encoding α-syn have been associated with familial forms of PD ^4^. While substantial progress has been made on understanding the normal function of α-syn ^5^ and its role in PD pathogenesis ^6^, comparatively less is known about the impact of post-translational modifications (PTMs) of α-syn and their role in PD pathogenesis.

PTMs are a series of modifications that influence protein localization, activity, and folding states. α-syn can undergo multiple different PTMs including but not limited to phosphorylation, glycation, acetylation, truncation, SUMOylation, and nitration ^7–10^. Phosphorylation of α-syn on serine 129 (pSyn) is the best characterized marker of Lewy bodies isolated from or imaged in patient post-mortem brain tissue. However, N-terminally acetylated α-syn (NAc-syn) is the most abundant endogenous form of α-syn *in vivo* ^11–13^. NAc-syn has been associated with a decreased propensity to form aggregates ^14–18^. This effect is believed to stem from the alteration of α-syn’s ability to form β-sheets thereby altering the structure of aggregates compared to non-acetylated α-syn. However, a more recent study found no differences in the structure of aggregates formed, instead pointing to a reduced propensity of acetylated α-syn aggregates to bind lipid membranes ^19^.

Lipid membrane binding is an important initial step for the internalization of α-syn aggerates into cells. Misfolded and aggregated α-syn is believed to promote the template-induced misfolding of native α-syn and spread from one cell to another in a prion-like manner ^20–22^. Multiple studies have demonstrated the spread of α-syn aggregates via different cell receptors present on cell membranes including heparin sulfate proteoglycans ^23–27^, but the effect of acetylation on cell surface receptor binding has not been studied extensively in model systems. Therefore, in addition to a reduction in aggregation propensity of α-syn, acetylation may alter the cell-to-cell propagation of α-syn aggregates, thereby limiting their toxic effects.

Thus, PTMs such as acetylation may play a crucial role in modifying the aggregation propensity, binding, and structural properties of α-syn, yet a clear elucidation of the cellular pathological effects of acetylation of α-syn using PD-relevant disease models is lacking. In this work, we report that N-terminal acetylation slowed α-syn aggregation kinetics *in vitro* via a seed amplification assay. Uptake of N-terminally acetylated α-syn pre-formed fibrils (NAc-PFFs) into immortalized cells and induced pluripotent stem cells (iPSC)-derived dopamine (DA) neurons was decreased compared to unmodified preformed fibrils (PFFs), possibly due to reduced binding to the cell surface. Finally, exposure to NAc-PFFs induced less seeding of endogenous α-syn than exposure to unmodified preformed fibrils (PFFs) in both iPSC-derived DA neurons and mouse brain. This also coincided with longer survival observed in mice injected with NAc-PFFs compared to PFFs.

## Materials and methods

### Generation and characterization of N-terminally acetylated α-synuclein preformed fibrils

Acetylation of the first methionine at the N-terminus of α-syn was performed as described in Rovere, Powers, Patel and Bartels ^28^. Briefly, *E.coli* were co-transformed with two plasmids, pET721a-α-syn (Addgene, #51486) and pTSara-NatB (a gift from Dr. Tim Bartels), which are responsible for expression of α-syn and the N-terminal acetylation enzyme NatB, respectively. Cultures were grown in LB+amp+cam media to OD ∼0.5-0.6 and induced with 0.2% L-arabinose 30 min before adding the second inducer (0.3mM isopropyl-β-D-thiogalactopyranoside) and further incubated for 4 hrs. NAc-α-syn in the clarified culture lysate after sonication was purified to homogeneity (>95% on SDS PAGE) by using a 5-ml HiTrap Q HP anion exchange column (GE Healthcare) and/or a 5-ml HiTrap Phenyl HP hydrophobic interaction column (GE Healthcare). N-terminal acetylation of α-syn was confirmed as by mass spectrometry (Figure 1A). Both α-syn and NAc-α-syn were tested to ensure endotoxin levels below 0.2 EU/mg prior to later steps. PFFs were generated by shaking 0.5-ml aliquot of unmodified or NAc-α-syn (5mg/ml) in 1.5-ml microtube at 1000rpm at 37°C for 7d. Additionally, both unmodified PFFs and NAc-PFFs were subjected to at least 40 cycles of 30s-On/30s-Off on a Bioruptor Pico sonication device (Diagenode). PFFs and NAc-PFFs were characterized using negative staining after sonication and visualized using a transmission electron microscope (Tecnai G2 Spirit Twin 120 kV Cryo-TEM) coupled to a camera (Gatan Ultrascan 4000 4 k × 4 k CCD Camera model 895) ^29^. Additional characterization of sonicated PFFs and NAc-PFFs was performed using dynamic light scattering (DLS). Images were analyzed with Fiji-ImageJ1.5 and GraphPad Prism 9 software.

**FIGURE 1.**
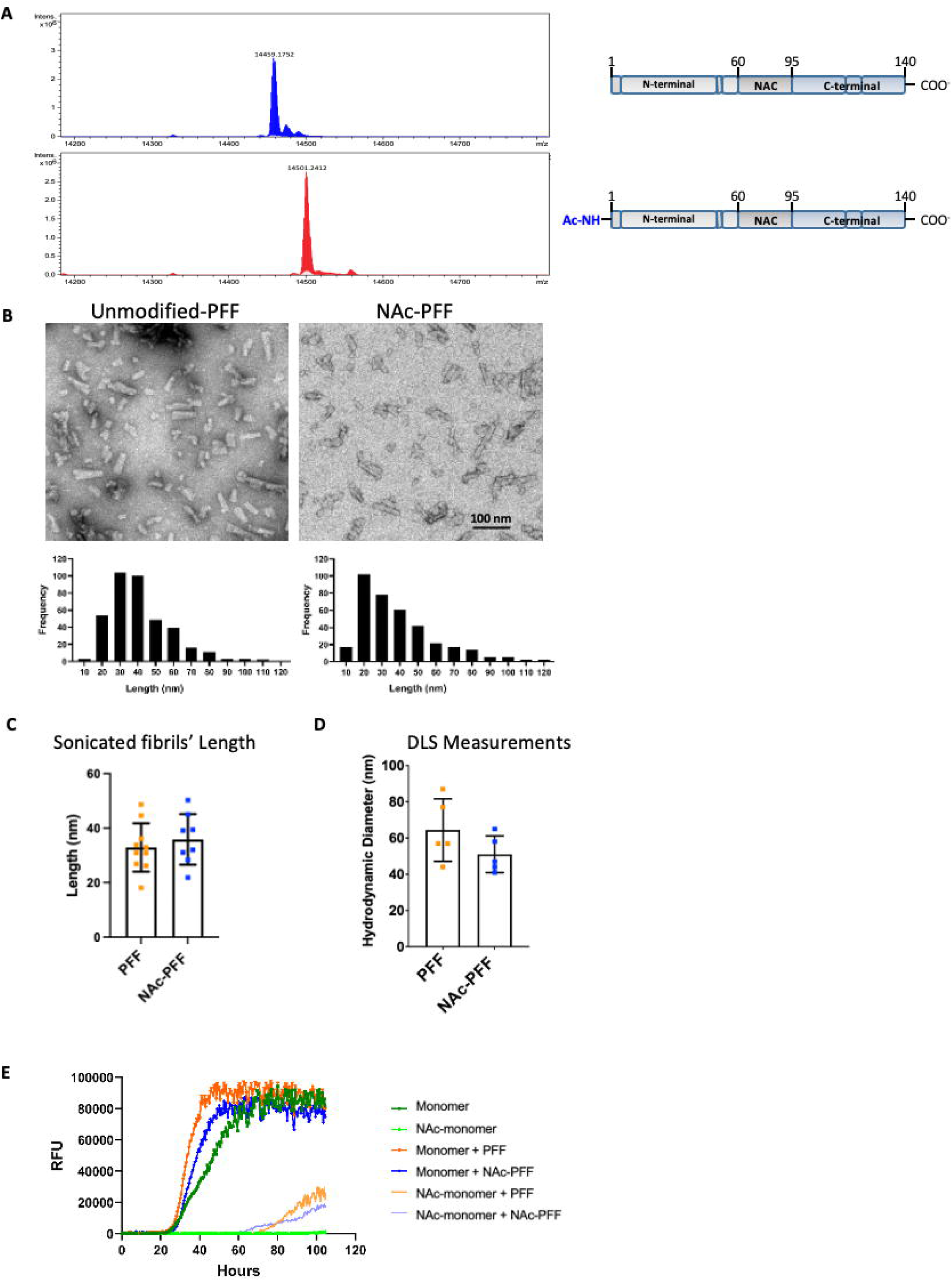
Characterization of PFFs vs NAc-PFFs. **A)** Mass spectrometry analyses of monomer construct used to make either PFF or NAc-PFF. Top spectrum reveals deconvoluted mass as expected from amino acid sequence for human WT α-syn sequence. Bottom spectrum shows full acetylation at the N-terminal of human WT α-syn sequence with a rightward shift (+42 Da). **B)** Electron micrograph images and characterization of length distributions for PFFs and NAc-PFFs post-sonication for a single batch each. **C)** Average length of fibrils after sonication of all batches of PFFs and NAc-PFFs. Statistical significance was assessed using unpaired t-tests. Bar graphs represent mean with error bars denoting ± SD. **D)** Hydrodynamic diameter measurements of 5 batches each of PFFs and NAc-PFFs post-sonication using dynamic light scattering. Statistical significance was assessed using unpaired t-tests. Bar graphs represent mean with error bars denoting ± SD. **E)** Seed amplification assay using α-syn with and without either PFF or N-terminal acetylation. Graphs display average of 4 wells.

### Seed amplification assay

Seed amplification assays were carried out as described previously with minor modifications ^30,31^. The base reaction mix was composed of 100 mM PIPES buffer pH 6.5, 300 mM NaCl, 10 uM ThT, and 0.002% SDS in filtered double distilled water. In all reactions, 0.1 mg/mL of monomer was used regardless of acetylation status, and 2 uL of 1:1000 diluted PFF was used to seed reactions (Final concentration in each well of 9 nM). Reactions were performed in quadruplicate technical replicates with a final volume of 100 uL per replicate in a 96-well plate format (corning). Plates were incubated in an Omega Fluostar plate reader (BMGLabtech) for 105 hours at 37°C. Plates were intermittently shaken (double orbital) with 1 min shaking and 29 min rest. Fluorescence readings were taken at 450 nm excitation and 480 nm emission every 30 min with a gain setting of 1,500.

### Generation of *SNCA* knockout and overexpressing RPE1 cells

For the generation of a *SNCA*-KO human retinal pigmented epithelial-1 cells (RPE1) cells, guide RNAs (gRNA1: GCCATGGATGTATTCATGAA and gRNA2: AAGCACCAAACTGACATTTG) were obtained from Integrated DNA Technologies (IDT) and dissolved in Nuclease-Free Duplex Buffer (IDT) according to the manufacturer’s instructions. The mixture was incubated at room temperature for 15 minutes to form the ribonucleoprotein (RNP) complex. Freshly isolated RPE1 cells (1.5 × 10⁵) were washed with PBS and resuspended in 75 μL of Nucleofector Solution mixed with Supplement 1 buffer (Lonza Nucleofector Kit V, Cat. No. VACA-1003), following the manufacturer’s protocol. The cells were then mixed with 25 μL of the prepared RNP complex by gentle pipetting and transferred to a nucleofection cuvette provided in the kit. Nucleofection was performed using the X-001 program on a Lonza Nucleofector device. Following nucleofection, the cells were transferred to a single well of a 12-well plate containing pre-warmed, antibiotic-free culture medium supplemented with 10% FBS and incubated at 37°C in a 5% CO₂ atmosphere. After 24 hours, the medium was replaced with fresh medium containing antibiotics. Genomic DNA was extracted from 0.5–1 × 10⁵ cells using QuickExtract DNA Extraction Solution (Lucigen, Cat. No. QE09050) according to the manufacturer’s protocol. DNA was processed in a thermal cycler with the following conditions: 65°C for 10 minutes, 68°C for 5 minutes, and 95°C for 5 minutes. Knockout of *SNCA* was confirmed by PCR using specific primers (Forward: TCCGTGGTTAGGTGGCTAGA, Reverse: CTGGAAAAGCAAACAGTCGCA), followed by sequencing and western blotting using an antibody against α-syn (BD Biosciences, Cat. No. 610787).

For wildtype (WT) *SNCA* expression, RPE1 endogenously express α-syn and so were used without any genetic modification. As for the generation of *SNCA* overexpression RPE1 cells, the coding sequence of WT α-syn was ordered as gBlock gene fragments (Integrated DNA Technologies). The insert was cloned into the NheI and BamHI sites of the doxycycline-inducible pCW57.1 plasmid (Addgene #41393, gift from David Root) using Gibson Assembly (New England Biolabs). PCR amplification was performed with Q5 DNA polymerase (NEB), and construct was sequence verified. Next, Lentiviral particles encoding WT α-syn were produced from the pCW57.1 vectors and used to transduce the RPE1 *SNCA*-KO cells above. Stable polyclonal cell populations were generated by selection with 15 µg/mL puromycin. Subsequently, upon addition of 2ug/ml of doxycycline (Sigma) overnight into the media, validation of *SNCA* mRNA and α-syn protein expression was conducted using RT-qPCR and Western blot analysis using primers targeting the *SNCA* gene or primary antibodies targeting α-syn protein, respectively.

RPE1 cells were grown and maintained at 37°C under a humidified atmosphere of 5% CO2. RPE1 cells were cultured in Dulbecco’s Modified Eagle’s Medium (Wisent) containing 4.5 g/l glucose, sodium pyruvate, supplemented with 10% heat inactivated fetal bovine serum (Wisent), 100 units (U)/ml penicillin, and 50 μg/ml streptomycin. Cells were passaged upon reaching approximately 70% - 90% confluence by rinsing in PBS followed by incubation in 0.05% Trypsin- EDTA (Wisent) at 37°C for approximately 5 min before propagation in a new cell culture dish or pelleted for experimental use. For propagation and passaging, trypsinized cells were re-suspended in fresh DMEM media to inactivate and dilute trypsin then counted using a Luna cell counter (Logos Biosystems) prior to plating at the desired cell density. For experimental use, trypsinized cells were resuspended in fresh media then split into 1.5 mL Eppendorf tubes or 15 mL falcon tubes depending on future use. Cells were pelleted by centrifuging at 1.2 rpm for 3 min, and the media was discarded via vacuum. Cell pellets were either stored at -20°C directly or re-suspended in cell lysis buffer and then stored at -20°C.

### iPSC culture and dopamine neuron differentiation

iPSCs from a PD patient with a triplication in the *SNCA* gene (and isogenic *SNCA* WT and *SNCA* KO iPSC lines generated by CRISPR-based gene editing from the same patient) were provided by Pr. Tilo Kunath from The University of Edinburgh ^32–34^. DA neuronal precursor cells (NPCs) and DA neurons were generated following previously established protocols ^35–37^. Briefly, iPSCs were dissociated with Gentle Cell dissociation reagent and transferred to uncoated flasks in NPC Induction Media to allow for embryoid bodies (EB’s) to form. EBs were cultured for 7 days and then transferred to polyornithine/ laminin coated flasks and grown for another 7 days in NPC induction media. To expand NPCs the EBs were then dissociated into small colonies by trituration in Gentle Cell dissociation media and replated as a monolayer on polyornithine/ laminin coated flasks. After reaching confluence, NPCs were harvested and frozen in FBS with 10% DMSO and stored in liquid nitrogen. To differentiate neurons, NPCs were thawed in NPC maintenance Media with Y-27632 (ROCK inhibitor) and plated on polyornithine/ laminin. NPCs were grown for 5-7 days until confluent. For final differentiation into DA neurons, NPCs were dissociated using Accutase and plated on polyornithine/ laminin in DA neuron differentiation media. DA neurons were maintained by exchanging 1/2 of the culture volume for fresh DA neuron differentiation media every 5-7 days. Neurons from every batch were assessed by immunofluorescence for expression of Map2 and TH. Only batches achieving at least 50% Map2/TH positivity after 4 weeks of differentiation were used for the experiments in this manuscript. Supplementary table 1 describes all media components used for iPSC culture and neuronal differentiation.

### PFF internalization, seeding, and uptake assays

RPE1 cells were seeded to 50% confluency in 96 well plates (Falcon), a day prior to treatment with 80nM of Alexafluor 488-labelled PFF or 488-labelled NAc-PFF. 24 hours post-treatment, cells were rinsed thrice with PBS, fixed with 4% PFA/PBS for 15min, and rinsed thrice in PBS. Cells were then processed for IF, before imaging and analysis. For DA neurons, NPCs were differentiated in 96-well plates (80,000 cells/ well) for 3 weeks and treated with vehicle (media), 80nM of Alexafluor 488-labelled PFF or 488-labelled NAc-PFF, for 6 or 24 hrs. Then, half of the media was removed and replaced with 8% PFA in PBS. The DA neurons were processed for IF followed by imaging and analysis. For seeding assays, DA neurons were treated with 300 nM unmodified PFF or NAc-PFF, followed by maintenance as described until desired timepoint (4- or 6-weeks post treatment).

### Immunofluorescence and high-content imaging

Cells were permeabilized for 10 min with 0.2% triton X-100 in PBS and blocked with 5% goat-serum and 0.02% triton X-100 in PBS. Antibodies used are described in Supplementary Table 2. High content imaging was performed on an Opera Phenix high-content confocal microscope (Perkin Elmer) and image analysis was performed using Columbus (Perkin Elmer). For DA neurons, data processing was then conducted using R studio as previously described ^35^. Nuclei were first identified by the Hoechst channel, and surrounding soma was identified as Map2-positive region. Relevant secondary stains were then quantified within this Map2-defined region. Single-cell data were then processed using a custom script in R studio to filter objects based on nuclear size, nuclear shape and Map2 staining intensity to identify only the neuronal cells for inclusion in subsequent analysis.

### Western Blot

Cultured cells were washed with PBS and lysed using RIPA buffer 0.1% SDS, 20mM Tris pH 7.4, 150mM NaCl, 1mM EDTA, 1mM EGTA, 1% NP-40, and 1% sodium deoxycholate, supplemented with phosphatase and protease inhibitors. Protein concentrations were determined via BCA, and samples were prepared in 6X Laemmli buffer supplemented with 0.1M DTT. 20 μg of total protein were loaded onto SDS-PAGE gel and separated using Biorad gel system. Gels were transferred onto nitrocellulose membranes (Biorad) and blocked for 1h in 5% skim milk made in 1X PBS with 0.1% Tween-20. For α-syn blots, membranes were fixed using 4% PFA and 0.1% glutaraldehyde for 30 min before blocking. Blocked membranes were incubated with primary antibody (Supplementary Table 2) at 4°C overnight. After, membranes were washed 3 times in 1X PBS with 0.1% Tween 20, and membranes were incubated with horseradish peroxidase (HRP)-conjugated secondary antibodies (Jackson ImmunoResearch) for 1h at room temperature. Protein detection was performed by capturing chemiluminescent signal on a ChemiDoc imaging system (Biorad) using Clarity/ Clarity max Western ECL Substrate (Biorad). Finally, blot signals were quantified using ImageJ.

### Intracerebral injection of PFFs in M83 transgenic mice

Three-month-old M83 hemizygous mice, overexpressing mutant human A53T α-syn under the PrP promoter ^38^ were bred in house, and maintained on a 12/12 h light/dark cycle at 22 °C ambient temperature with unlimited access to food and water. Housing, breeding, and procedures were performed according to the Canadian Council on Animal Care and were approved by the McGill University Animal Care Committee. Prior to intracerebral injection, M83 mice (females and males) were housed in individual plastic cages; and prior to the craniotomy, a dose of 20 mg/kg carprofen and 250 mg/kg bupivacaine were administered subcutaneously. M83 mice were anesthetized with 2% isoflurane and stereotaxically injected with PBS, PFFs or NAc-PFFs (12.5 μg per brain), in the right dorsal striatum using a 5 µl Hamilton syringe with a 33-gauge needle (Luk et al., 2012). All mice were injected the same day and were divided into two groups, endpoint (n= 20) and survival (n= 20). Mice were weighed before and, at regular intervals, after injection. Humane endpoint was defined as the point in time when motor impairment became debilitating (severe loss of balance preventing proper food and water consumption), or if the mice lost more than 20% of their starting body weight. For the endpoint group, the mice were euthanized when the first mouse reached the humane endpoint. For the survival group, each mouse was euthanized when it reached its individual humane endpoint. Mice were anesthetized with 2% isoflurane and were perfused with PBS followed by 10% formalin. The brains were removed and postfixed with 10% formalin for 24h at 4°C. Brains were then incubated in 70% ethanol for a week before processing for paraffin embedding.

### Immunohistochemistry and immunofluorescence of M83 mice brains

Coronal brain sections were cut with a paraffin microtome at 5 μm thickness. Tissue sections were incubated in citrate buffer (pH 6.0) for 10 min, rinsed with Tris-buffered saline containing 0.1% Tween-20 (TBST), and incubated in 3% oxidase peroxidase for 15 min. Subsequently, the sections were blocked with 10% normal goat serum in TBST for 30 min at room temperature and incubated with anti-pSyn antibody (abcam, ab184674) overnight at 4°C. The next day, sections were incubated with horseradish peroxidase (HRP)-conjugated secondary antibodies (Jackson ImmunoResearch) for 30 min. After washing with PBS, cells were analyzed. The peroxidase reaction product was visualized as a brown precipitate by incubating the tissue with the DAB substrate kit (8059, cell signal technology). Sections were additionally incubated in hematoxylin to stain for nuclei. Three coronal sections containing the substantia nigra (minimal distance between sections 40 μm), were examined by a bright-field microscope (Olympus DP-21SAL coupled to a digital camera DP21/DP26). Then the images were analyzed using Fiji-ImageJ 1.53 software. Macros for Fiji-ImageJ were written in Jython, using basic ImageJ functions. We defined each individual spot labeled with pSyn as a positive particle for pSyn. Cell particles were defined by perimeter, area, and size.

For immunofluorescence staining, coronal brain sections were blocked in 10% normal goat serum in TBST for 1 hour at room temperature and incubated with anti-pSyn antibody and anti-TH overnight at 4°C. After rinsing with PBS, the tissue sections were incubated with anti-mouse Alexa fluor-488, anti-rabbit Alexa-Fluor-555 and DAPI for 2 hrs at room temperature. Coronal sections were examined using a Zeiss Axio Observer Z1 microscope. Two midbrain coronal sections (minimal distance between sections 40 μm) were analyzed using Fiji-ImageJ 1.53 software. Macros for Fiji-ImageJ were written in Jython, using basic ImageJ functions.

### Statistical Analysis

All statistical analyses were conducted in GraphPad Prism9 software. For experiments involving retinal pigment epithelial cells (RPE1), a minimum of three replicates were utilized for each of the three experiments (n=3) performed for both the uptake assays and binding assays. For experiments in DA neurons, biological replicates were defined as separate individual differentiations of banked NPCs to neurons (n= 3). In the graphs, each point represents an average of technical replicates (between 2-3) for each batch of PFF (unmodified PFFs or NAc-PFFs) per experiment. Statistical comparisons were performed using unpaired t-tests, One- Way ANOVA or Two-Way ANOVA followed by Tukey-corrected tests or Dunnett’s multiple comparison test where applicable.

## Results

### Biochemical and structural properties of unmodified and N-terminally acetylated α-synuclein preformed fibrils

Both unmodified and N-terminally acetylated wildtype human α-syn were produced in *E. Coli* and purified to homogeneity. Subsequent analysis by mass spectrometry showed complete N-terminal acetylation (Fig. 1A). PFFs or NAc-PFFs were then generated from their corresponding monomers in separate batches. After sonication, the length of both types of PFFs appeared similar, and this was quantified both using electron microscopy and dynamic light scattering measurements (Fig. 1B-D). We then sought to characterize the kinetics of aggregate formation using seed amplification assay. ThioflavinT selectively fluoresces when it binds to β-sheet-rich regions in protein aggregates monitored in real-time using a plate reader ^39^. We observed that acetylation of either the monomer, PFFs, or both lead to a markedly longer lag phase and lower max RFU after 100 hours (Fig. 1E). These effects of acetylation on the kinetics of monomeric α-syn are in line with previous literature ^17,18,40^.

### N-terminally acetylated α-synuclein preformed fibrils are internalized less efficiently by RPE1 cells

Transmission of α-syn has been widely reported to occur in a prion-like manner in patients and animal models ^41–45^, and many mechanisms have been suggested for this process including endocytic vesicles, receptor-mediated uptake, and phagocytosis ^46–50^. PTMs such as acetylation have significant impact on α-syn’s stability, localization, and kinetics ^16,51^. Therefore, we sought to evaluate and monitor the uptake of unmodified and NAc-PFFs using RPE1 cells that express varying levels of α-syn (*SNCA*-WT, *SNCA*-KO, and *SNCA*-overexpression) (Fig. 2A). The purpose of this is two-fold: understand the impact of acetylation on uptake of α-syn, and the effect to which endogenous levels of α-syn may affect this process. We found that treatment with 15nM of 488nm-PFFs (either unmodified- or NAc-) for 1hr showed an approximately 50% reduction in uptake of NAc-PFFs compared to unmodified PFFs (Fig. 2B-C), regardless of the *SNCA* genotype of the cells. Similar results were observed with a longer incubation period of 24hrs (Fig. 2D). These findings indicate that, in RPE1 cells, the initial internalization and accumulation steps of both unmodified- and NAc-PFFs are independent of endogenous α-syn levels but that NAc-PFF uptake is reduced compared to unmodified PFFs.

**FIGURE 2.**
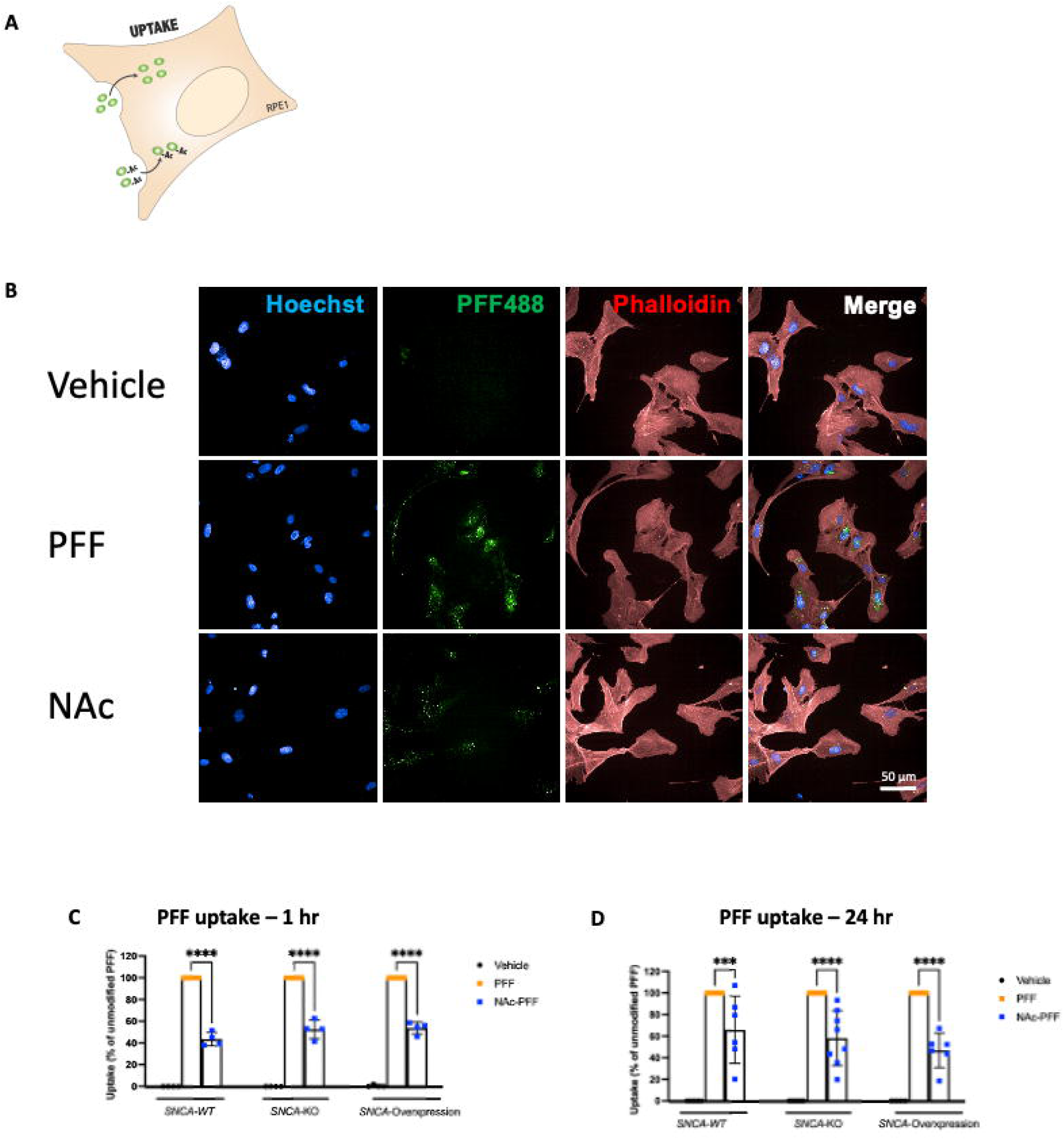
Fibril uptake is reduced in RPE1 cells with NAc-PFF compared to unmodified PFF. **A)** Schematic for experimental setup utilizing RPE1 cells to assess uptake of 488-AlexaFluor labelled PFF vs NAc-PFF fibrils. **B)** Representative immunofluorescence images of *SNCA*-WT RPE1 uptake assay at 1 hr timepoint. **C)** Quantification of labelled PFF uptake in RPE1 cells with different levels of endogenous *SNCA* expression (WT, KO, and overexpression) at 1 hr and 24 hr. Data was first normalized by subtracting vehicle for each cell line, and 488-PFF was set to 100% uptake. 488-NAc-PFF was expressed as a percent compared to 488-PFF. Statistical significance was assessed using two-way ANOVA followed by Dunnett’s multiple comparison test. Bar graphs represent mean with error bars denoting ± SD. Significance levels are depicted in figure legends (“***” indicates p≤0.0005, “****” indicates p<0.0001).

### Cleavage of extracellular heparan sulfate proteoglycans reduces cell surface binding of unmodified and N-terminally acetylated α-synuclein preformed fibrils in RPE1 cells

To assess the mechanism of initial binding of unmodified- and NAc-PFF to the cell surface, labelled PFFs were incubated with RPE1 expressing endogenous levels of α-syn (*SNCA-*WT) for 20 min on ice to inhibit any active endocytic mechanisms involved in PFF internalization ^46^ (Fig. 3A). We observed over 70% decrease in the amount of NAc-PFF bound to the cell surface compared to unmodified PFF (Fig. 3B-C). Previously, we and other groups have shown that heparan sulfate proteoglycans (HSPGs) facilitate the uptake of α-syn fibrils ^23,24^. To this end, we pretreated *SNCA*-WT RPE1 cells with a cocktail of heparin lyases enzymes (heparinase) for 1 hr. This cocktail, containing Heparinase I and III, provided us an almost complete and selective cleavage of the heparin and heparan sulfate chains confirmed through immunofluorescence staining. By quantifying HSPG area per cell, we observe 1h pre-treatment to be sufficient in abolishing the HSPG fluorescence on the cell surface compared to untreated cells (Fig. 3B and D). Following this, we observed an approximate 80% reduction in unmodified PFF intensity per cell after Heparinase treatment (Fig. 3B-C), indicating a decrease in binding and internalization. Interestingly, we also observed an almost complete loss of NAc-PFF binding to the cell surface after heparinase treatment (Fig. 3B-C). Taken together, these findings indicate that NAc-PFF has reduced binding efficiency compared to PFF.

**FIGURE 3.**
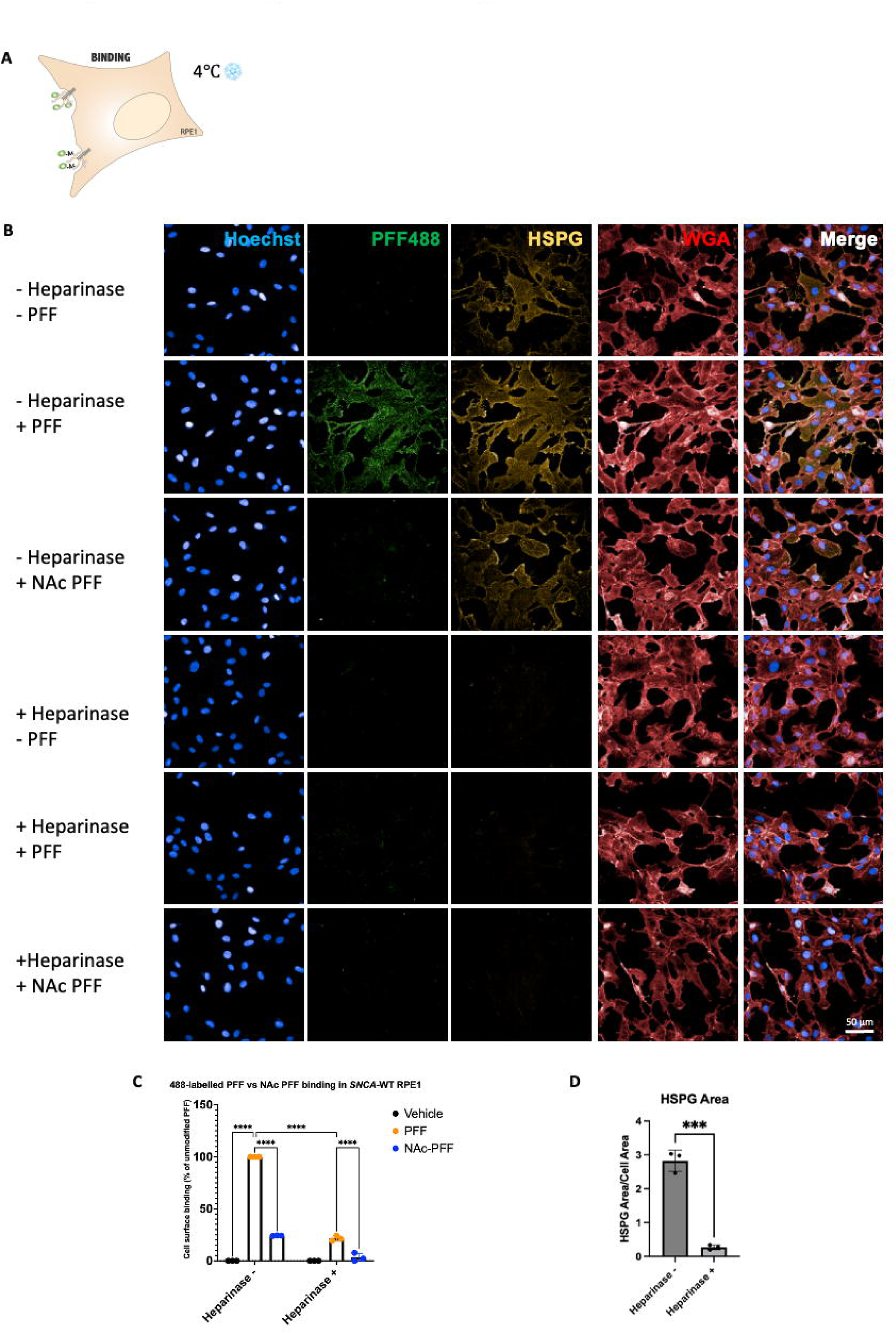
Heparinase treatment abrogates PFF and NAc-PFF binding in RPE1 cells. **A)** Schematic for experimental setup utilizing RPE1 cells to assess binding affinity of 488-AlexaFluor labelled PFF vs NAc-PFF fibrils and hypothesized interaction via HSPGs. **B)** Representative immunofluorescence images of *SNCA*-WT RPE1 binding to 488-PFF vs 488-NAc-PFF with and without heparinase. **C)** Quantification of labelled PFF binding in *SNCA*-WT RPE1. Data was first normalized by subtracting vehicle condition to remove background fluorescence. Then, percent binding for 488-PFF without heparinase was set to 100%. All other conditions were expressed as percent cell surface binding compared to the PFF condition. Statistical significance was assessed using two-way ANOVA followed by Tukey’s multiple comparison test. Bar graphs represent mean with error bars denoting ± SD. **D)** Quantification of HSPG area per cell area with and without heparinase. Statistical significance was assed using unpaired t-test. Bar graphs represent mean with error bars denoting ± SD. Significance levels are depicted in figure legends (“***” indicates p≤0.0005, “****” indicates p<0.0001).

### N-terminally acetylated α-synuclein preformed fibrils are internalized less efficiently by iPSC-derived dopamine neurons

To test whether NAc-PFF uptake is also reduced in human patient-derived dopamine DA neurons, we used iPSCs generated from a PD patient with a triplication of the *SNCA* gene (4 copies of the *SNCA* gene; *SNCA*-Triplication), as well as isogenic CRISPR edited iPSC lines with either 2 copies (*SNCA*-WT) or zero copies (*SNCA*-KO) of the *SNCA* gene ^32–35^. Similar to the approach in RPE1 cells (Fig. 2), we tracked the uptake of 488nm-PFFs (unmodified- or NAc-) over time (Fig. 4A). We observed that, in DA neurons from all three isogenic iPSC lines, NAc-PFFs were internalized less than unmodified PFFs (Fig. 4B-C) independent of endogenous α-syn expression. This difference was also observed at 24h but did not meet the threshold for statical significance (Fig. 4B-C) potentially indicating a levelling-off effect at this time point.

**FIGURE 4.**
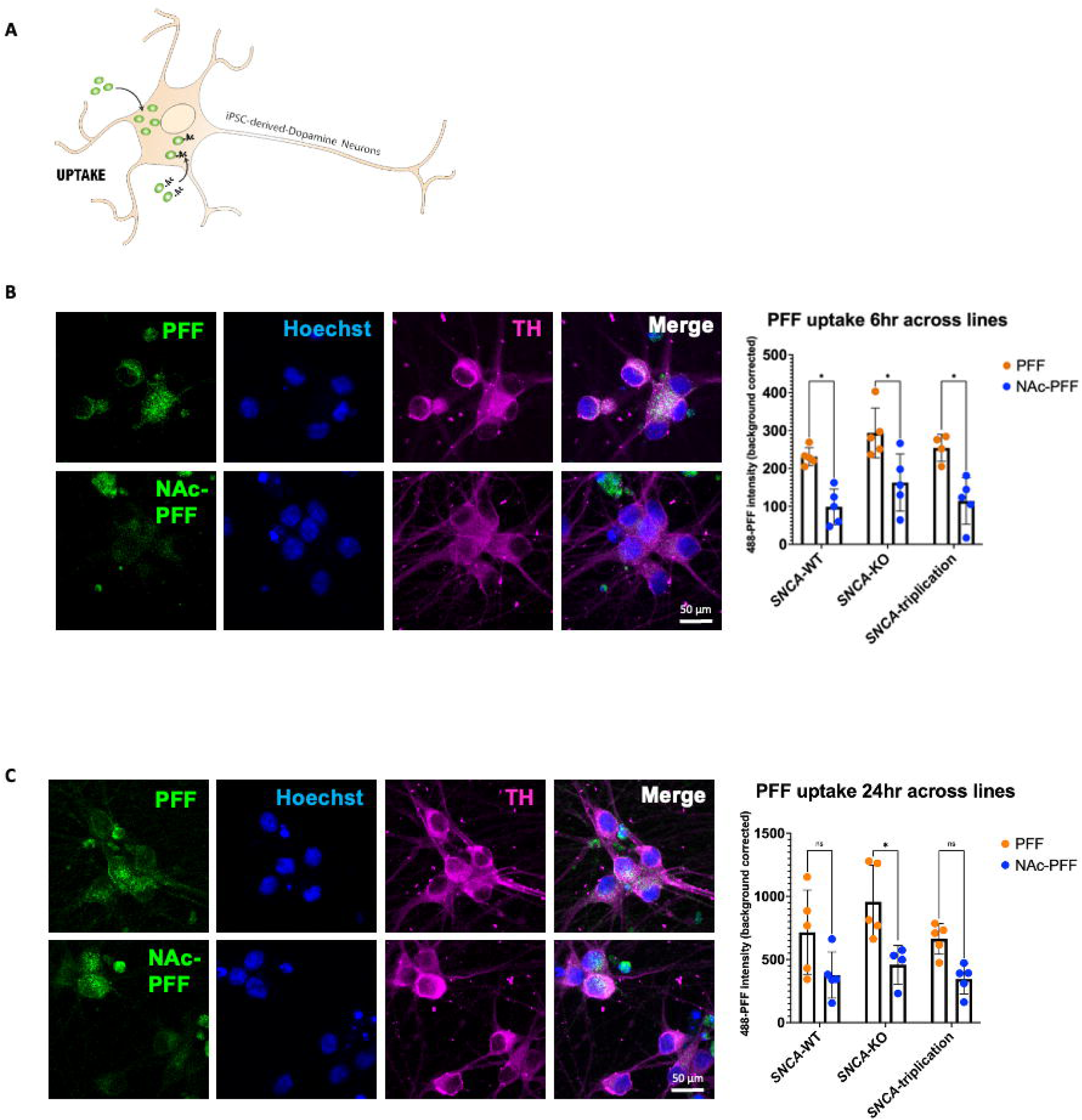
NAc-PFFs exhibit reduced uptake in iPSC-derived dopamine neurons regardless of endogenous *SNCA* expression levels. **A)** Schematic for experimental setup utilizing DA neurons assessing 488-labelled PFF vs NAc-PFF uptake. **B)** Representative immunofluorescence images of *SNCA*-WT iPSC-derived DA neurons uptake of 488-PFF vs 488-NAc-PFF at 6 hr. **C)** Representative immunofluorescence images of *SNCA*-WT iPSC-derived DA neurons uptake of 488-PFF vs 488-NAc-PFF at 24 hr. Quantification was performed for all three levels of α-syn expression in neurons (*SNCA*-WT, *SNCA*-KO, and *SNCA*-Triplication). All data was background corrected for each line by subtracting vehicle treatment condition. Statistical significance was assessed using two-way ANOVA followed by Tukey’s multiple comparison test. Bar graphs represent mean with error bars denoting ± SD. Significance levels are depicted in figure legends (“*” indicates p<0.05).

### N-terminally acetylated α-synuclein preformed fibrils induce less α-synuclein seeding in dopamine neurons

We next sought to determine if N-terminal acetylation also affected the capacity of exogenous PFFs to seed endogenous α-syn aggregates in DA neurons (Fig. 5A). As most intracellular α-syn inclusions, including Lewy bodies in the brains of PD patients, contain α-syn that is phosphorylated on serine 129, we used pSyn staining as a surrogate for α-syn seeding. We first examined the total α-syn in Map2-TH double-positive neurons and confirmed that *SNCA*-KO lacked α-syn expression while *SNCA*-triplication had higher levels of total α-syn than the isogenic *SNCA*-WT (Supplementary Fig. 1). Next, *SNCA*-Triplication, *SNCA*-WT and *SNCA*-KO DA neurons were treated with 300 nM of unlabeled unmodified- or NAc-PFFs and maintained in culture for 4 and 6 weeks prior to fixation and analysis (Fig. 5B-C). At both 4 and 6 weeks timepoints, consistent with the well-established dependence of seeding on elevated endogenous α-syn levels ^33,52^, we observed a statistically significant increase in seeding in the *SNCA*-Triplication line. More importantly, at both time points, unmodified-PFFs induced greater α-syn seeding than NAc-PFF. Together, these results show that, compared to unmodified α-syn PFFs, N-terminally acetylated PFFs induce less seeding in DA neurons, consistent with their propensity for reduced binding, uptake and *in vitro* fibrillization.

**FIGURE 5.**
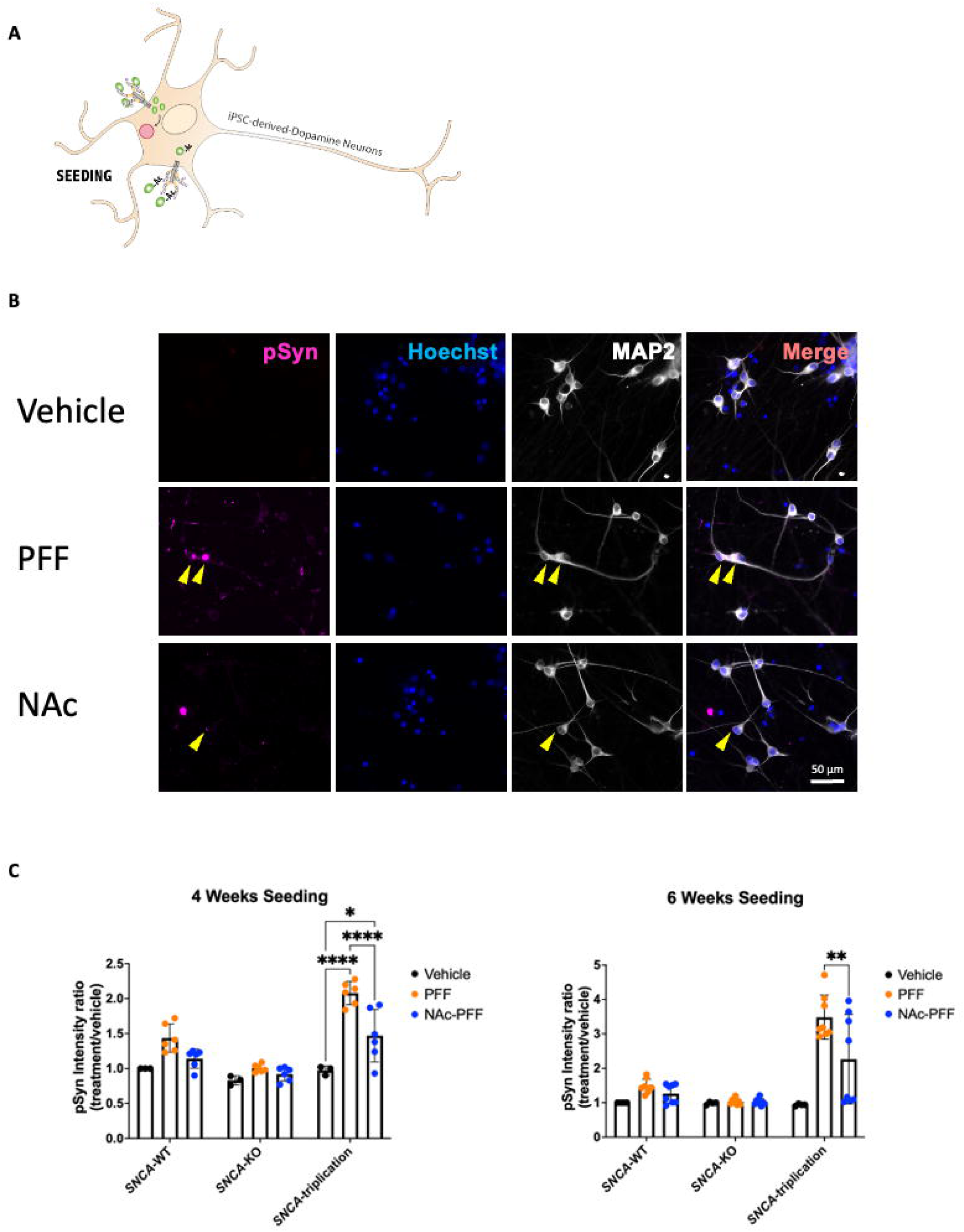
NAc-PFFs exhibit reduced seeding efficiency in iPSC-derived dopamine neurons. **A)** Schematic for experimental setup utilizing DA neurons assessing PFF vs NAc-PFF seeding activity. **B)** Representative immunofluorescence images of *SNCA*-WT iPSC-derived DA neurons treated with PFFs or NAc-PFFs for 4 weeks. Yellow arrowheads indicate intracellular pSyn inclusions used for quantification of seeding. **C)** Quantification of seeding efficiency measured by pSyn staining intensity in all three lines of iPSC-derived DA neurons (*SNCA*-WT, *SNCA*-KO, and *SNCA*-Triplication) for 4 weeks and 6 weeks. Data is expressed as pSyn intensity ratio for each treatment condition over baseline (*SNCA*-WT, vehicle treated). Statistical significance was assessed using two-way ANOVA followed by Tukey’s multiple comparison test. Bar graphs represent mean with error bars denoting ± SD. Significance levels are depicted in figure legends (“*” indicates p<0.05, “**” indicates p≤0.005, “****” indicates p<0.0001).

### M83 α-synuclein hemizygous transgenic mice injected with N-terminally acetylated α-synuclein preformed fibrils survive longer than mice injected with unmodified preformed fibrils

To address the role of the N-terminal acetylation of α-syn *in vivo*, we used a well-characterized mouse model of spreading synucleinopathy ^38^. All M83 hemizygous mice were injected the same day with PBS (control), unmodified PFFs or NAc-PFFs, in the right dorsal striatum (n= 40), and we divided the mice into two groups – survival and endpoint experiments. Fig. 6A summarizes the experimental timeline. For the survival experiment, we allowed each mouse to reach its individual humane endpoint before euthanizing it; all the mice injected with unmodified-PFFs developed symptoms and reached their humane endpoint before the first mouse injected with NAc-PFF became symptomatic (Fig. 6B). For the endpoint experiment, all mice were euthanized at the same time point once the first mouse reached the humane endpoint. This first mouse was part of the group injected with unmodified-PFFs. In both groups NAc- and unmodified-PFF injected mice, pSyn positive profiles were found in cell soma, dendritic branches and axons (Fig. 6C) of the substantia nigra (SN) whereas no pSyn staining was observed in the control mice (PBS-injected group). Importantly, pSyn staining was significantly lower in the substantia nigra (Fig. 6D) and pons (Fig. 6E) of mice injected with NAc-PFFs compared to those injected with unmodified PFFs. In contrast, no such reduction was observed in pSyn staining adjacent to the site of NAc-PFF injections in the right striatum (Fig. 6F), with even a slight trend towards increased pSyn staining in this region when mice were injected with NAc-PFF. We also used immunofluorescence to determine if the DA neurons, positive for TH, contained pSyn in the SN. In agreement with our *in vitro* studies, the percentage of TH-positive neurons containing pSyn appeared higher in the mice injected with unmodified-PFFs compared to the NAc-PFF, although this did not reach statistical significance (Fig. 6G-H). Overall, our results suggest that mice injected with NAc-PFF survive longer and that NAc-PFF propagation in the brain is slower and less extensive than for unmodified-PFFs.

**FIGURE 6.**
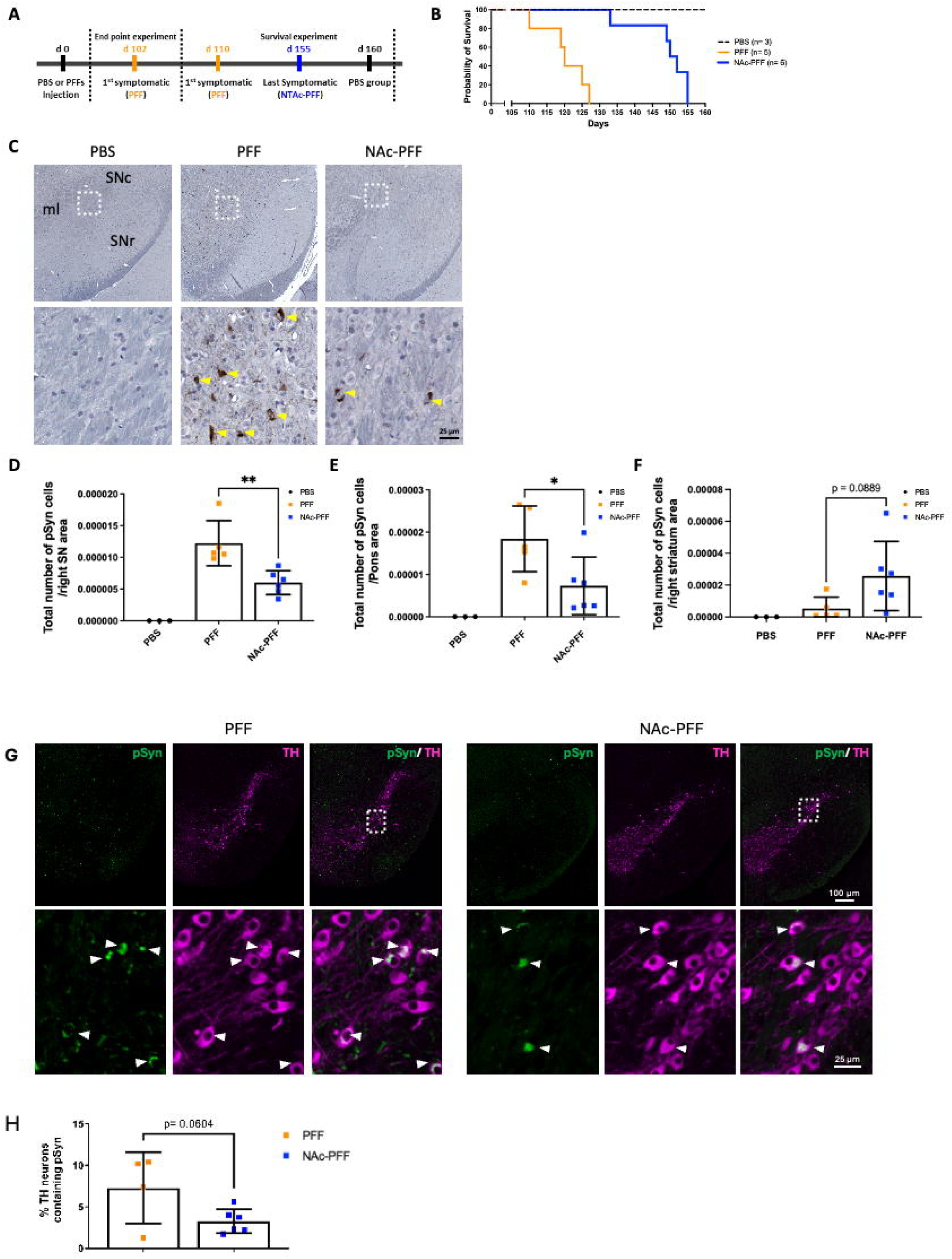
PFF injections lead to shorter survival and higher levels of phosphorylated synuclein than NAc-PFFs *in vivo*. **A)** Experimental timeline of mice models indicating end point and survival paradigms. **B)** Survival curves for mice injected with PBS, PFF, or NAc-PFF in the survival paradigm. **C)** Representative immunohistochemistry staining pSyn within the substantia nigra of mice injected with PBS or PFFs (unmodified or N-terminally acetylated). Yellow arrows indicate phosphorylated α-syn inclusions identified**. D)** Quantification of phosphorylated α-syn inclusions in the substantia nigra, **E)** pons and **F)** right striatum (Site of PFF injection). Statistical significance was assessed using one-way ANOVA. Bar graphs represent mean with error bars denoting ± SD. (“*” indicates p<0.05, “**” indicates p≤0.005) **G)** Representative images showing colocalization of TH and pSyn within neurons of the substantia nigra injected with either PFFs (n= 4) or NAc-PFFs (n= 6). White arrows indicate pSyn inclusions co-localized to TH+ neurons used for quantification. **H)** Quantification of percent TH neurons that show colocalization with pSyn in the substantia nigra. Statistical significance was assessed using unpaired t-test. Bar graphs represent mean with error bars denoting ± SD.

## Discussion

In the present study, we provide important *in vitro* and *in vivo* evidence supporting that PFFs generated from N-terminally acetylated monomeric α-syn have a lower propensity for fibrillization, binding to the cell surface, internalization and cell-to-cell transmission. We demonstrate in RPE1 cells and iPSC-derived DA neurons that NAc-PFFs are internalized less readily than unmodified PFFs, regardless of endogenous α-syn expression levels. Furthermore, we propose that this is likely due to reduced binding to the cell surface via HSPG-modified receptors. Several receptors have been identified to bind α-syn, including HSPGs, LAG3, and neurexin 1β ^23,27,53,54^. It is described that unmodified-PFF are internalized using HSPG-modified receptors whereas NAc-PFFs bind N-linked glycans and neurexin 1β ^53^. However, our findings using heparinase indicate that HSPG are also involved, at least in part, in NAc-PFFs binding. Previous studies have shown that lipids and lipid-binding partners can be an important driver for the aggregation of α-syn *in vitro* ^19,55–57^. As such, lower binding propensity to lipid membranes via acetylation may not only impact α-syn’s endogenous function but also alter the cellular pathology involved in PD which can be of focus for future studies.

We find that N-terminal acetylation has a significant effect on the pathological impact of PFFs. We demonstrate that iPSC-derived DA neurons incubated with NAc-PFFs developed less pathological pSyn-positive inclusions than those treated with unmodified PFFs. This could be explained in several ways. First, reduced binding to the cell surface as shown in RPE1 cells could lead to less uptake and subsequent reduced exposure to fibrils that seed pathology within cells. Second, our *in vitro* data (ThT assay) indicate that NAc-PFFs have a lower propensity for α-syn monomer recruitment, which could explain the comparatively lower seeding of endogenous α-syn in cells. The effect of N-terminal acetylation on α-syn aggregation properties is controversial. Some studies have shown that NAc has little or no effect on α-syn aggregation ^58^; whereas other studies have shown slower α-syn aggregation and fibril formation ^15–17,40,57^ which is consistent with our results. Yet another study by Birol, Wojcik, Miranker and Rhoades ^53^, showed higher seeding efficiency in primary neurons treated with acetylated PFFs, which contrasts with our findings. This difference may be due to differences in the methods of PFF generation, as it has been previously shown that PFFs generated in the presence of salt can lead to distinct, more pathologically potent strains of PFFs than those formed without salt ^59^. Additionally, differences in neuronal culture models might also explain these diverging results.

Finally, we demonstrate in a well-characterized *in vivo* mouse model that injection of NAc-PFFs lead to slower spreading of α-syn pathology as shown by increased survival and decreased presence of pSyn inclusions in disease-relevant brain regions. PFFs have been extensively used to induce toxicity in mouse models ^60,61^. We hypothesize that the attenuated spreading of NAc-PFF-induced pathology throughout the brain is due to, at least in part, to their reduced propensity for cell surface binding, internalization and seeding. We can even speculate that the trend towards higher amounts of pSyn observed at the NAc-PFF injection sites compared to the unmodified-PFF injection sites reflects a lack of spreading and propagation N-terminally acetylated form.

PTMs can have significant impact on the structural, biochemical, and physiological properties of proteins. While PTMs of α-syn have been well-described, much of their physiological and pathological functions remain to be elucidated. In this work, for the first time, we demonstrate that PFFs generated from N-terminally acetylated α-syn exhibit fundamentally altered cell surface binding properties and reduced internalization, seeding and cell-to-cell spreading both *in vitro* and *in vivo.* Thus, more broadly, our work points to the possibility that an imbalance or dysregulation of N-terminal acetylation may impact α-syn pathology or the rate of disease progression of synucleinopathies such as PD.

## Data availability

The authors confirm that the data supporting the findings of this study are available within the article [and/or] its supplementary material.

## Acknowledgements

We thank Professor Tim Bartels for the pTSara-NatB plasmid. We thank Tilo Kunath for the patient-derived SNCA triplication iPSC line and for the CRISPR-corrected isogenic *SNCA* WT and *SNCA* KO iPSC lines generated from the same patient.

## Supplementary material

Supplementary material is provided online.

---

**Figure S1. Synuclein expression in iPSc-derived dopamine neurons.** Immunofluorescence quantification of α-syn expression levels in SNCA-WT, SNCA-KO, and SNCA-Triplication iPSC-derived DA neurons. Data was normalized by subtracting KO signal to account for any background. Statistical significance was assessed using two-way ANOVA followed by Tukey’s multiple comparison test. Bar graphs represent mean with error bars denoting ± SD. (“****” indicates p<0.0001)

**Supplementary Table 1.** List of reagents used for iPSC and DA neuronal induction, differentiation and maintenance.

**Supplementary Table 2.** List of antibodies and their dilutions used in the current study for immunofluorescence, immunohistochemistry and Western blot.

