## Supplemental for "N-terminal acetylation reduces α-synuclein pathology in models of Parkinson’s disease"

### Slide 1
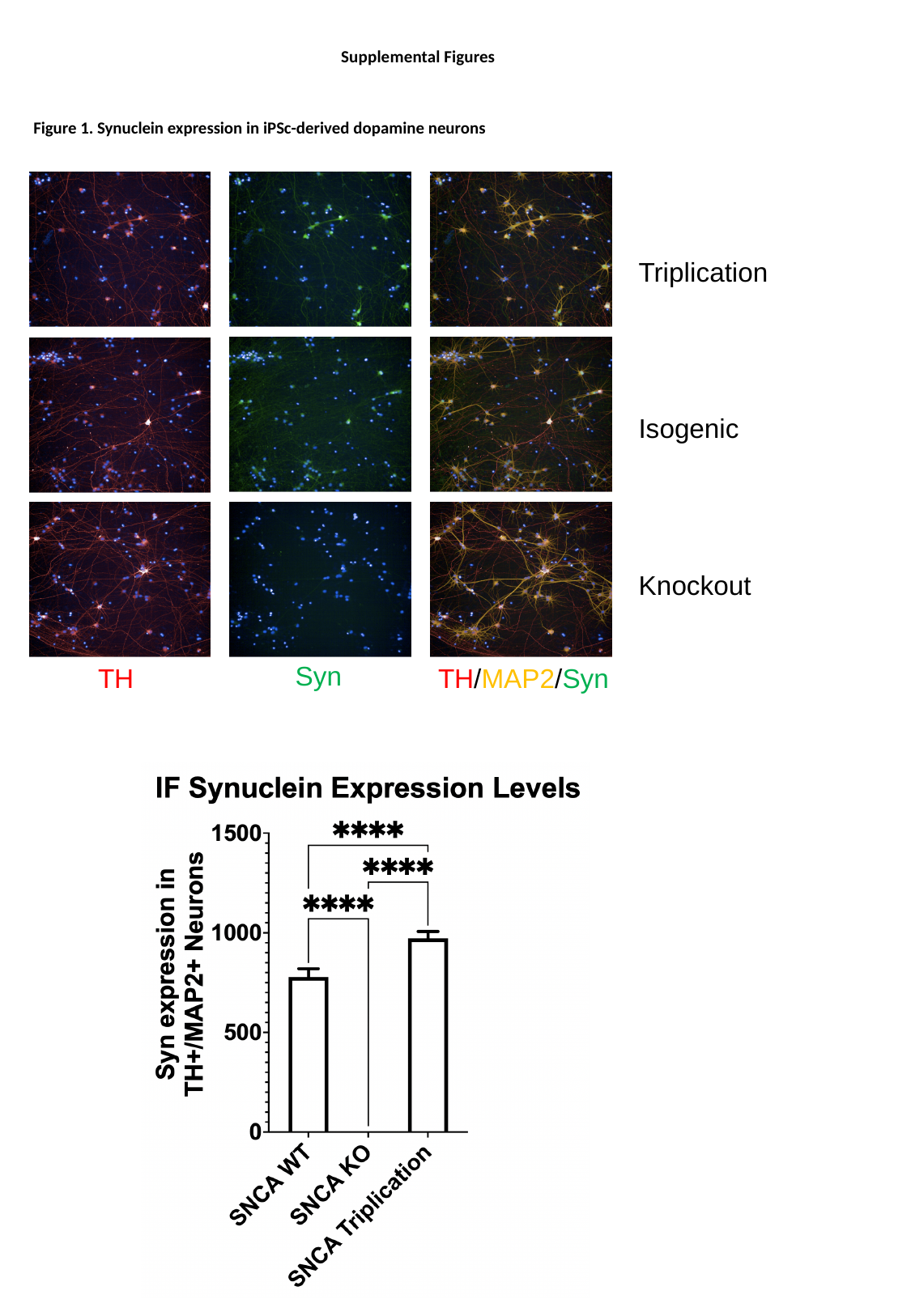

Supplemental Figures
Figure 1. Synuclein expression in iPSc-derived dopamine neurons
Triplication
Isogenic
Knockout
Syn
TH
TH/MAP2/Syn

### Slide 2
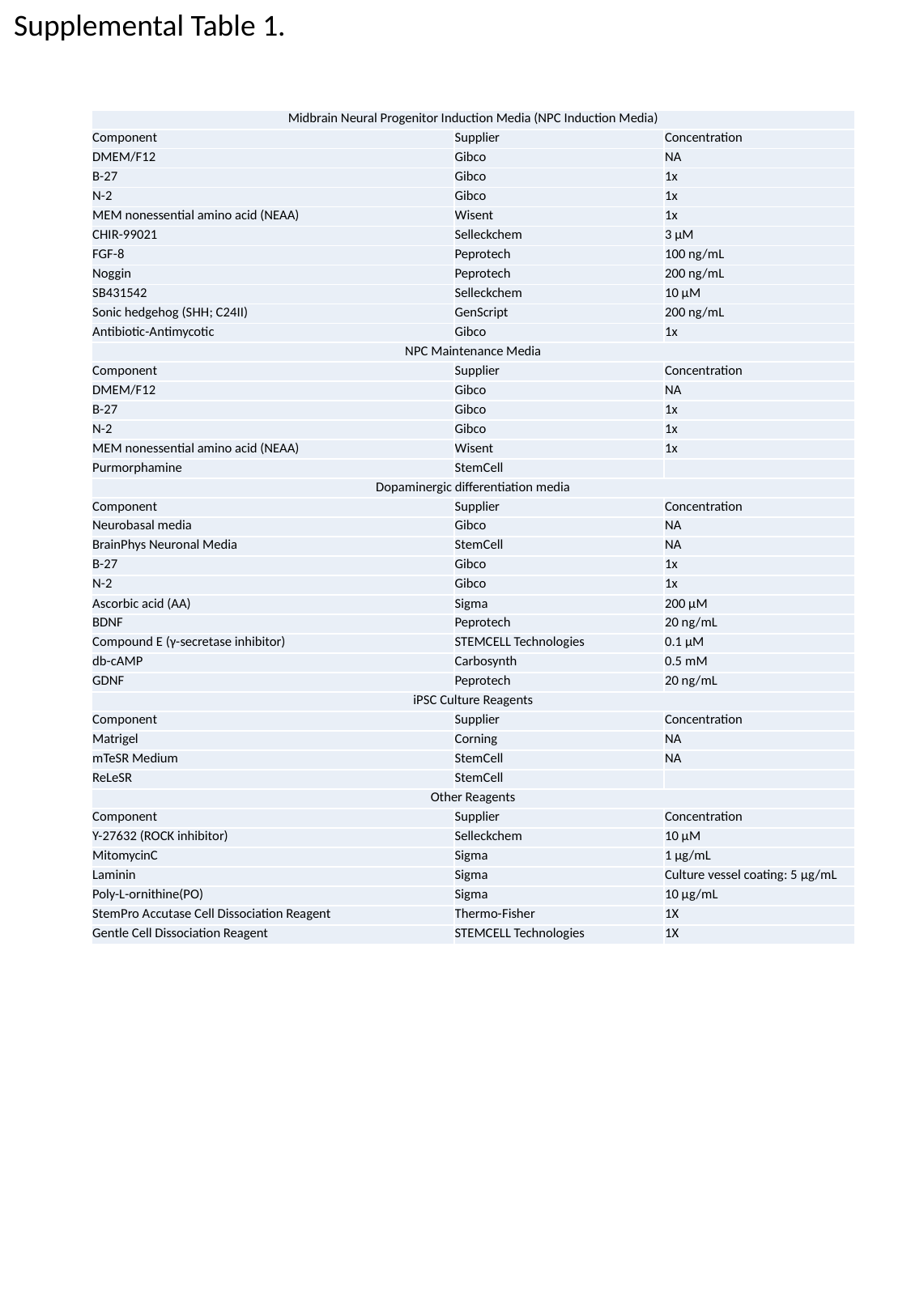

Supplemental Table 1.
| Midbrain Neural Progenitor Induction Media (NPC Induction Media) | | |
| --- | --- | --- |
| Component | Supplier | Concentration |
| DMEM/F12 | Gibco | NA |
| B-27 | Gibco | 1x |
| N-2 | Gibco | 1x |
| MEM nonessential amino acid (NEAA) | Wisent | 1x |
| CHIR-99021 | Selleckchem | 3 μM |
| FGF-8 | Peprotech | 100 ng/mL |
| Noggin | Peprotech | 200 ng/mL |
| SB431542 | Selleckchem | 10 μM |
| Sonic hedgehog (SHH; C24II) | GenScript | 200 ng/mL |
| Antibiotic-Antimycotic | Gibco | 1x |
| NPC Maintenance Media | | |
| Component | Supplier | Concentration |
| DMEM/F12 | Gibco | NA |
| B-27 | Gibco | 1x |
| N-2 | Gibco | 1x |
| MEM nonessential amino acid (NEAA) | Wisent | 1x |
| Purmorphamine | StemCell | |
| Dopaminergic differentiation media | | |
| Component | Supplier | Concentration |
| Neurobasal media | Gibco | NA |
| BrainPhys Neuronal Media | StemCell | NA |
| B-27 | Gibco | 1x |
| N-2 | Gibco | 1x |
| Ascorbic acid (AA) | Sigma | 200 μM |
| BDNF | Peprotech | 20 ng/mL |
| Compound E (γ-secretase inhibitor) | STEMCELL Technologies | 0.1 μM |
| db-cAMP | Carbosynth | 0.5 mM |
| GDNF | Peprotech | 20 ng/mL |
| iPSC Culture Reagents | | |
| Component | Supplier | Concentration |
| Matrigel | Corning | NA |
| mTeSR Medium | StemCell | NA |
| ReLeSR | StemCell | |
| Other Reagents | | |
| Component | Supplier | Concentration |
| Y-27632 (ROCK inhibitor) | Selleckchem | 10 μM |
| MitomycinC | Sigma | 1 μg/mL |
| Laminin | Sigma | Culture vessel coating: 5 μg/mL |
| Poly-L-ornithine(PO) | Sigma | 10 μg/mL |
| StemPro Accutase Cell Dissociation Reagent | Thermo-Fisher | 1X |
| Gentle Cell Dissociation Reagent | STEMCELL Technologies | 1X |

### Slide 3
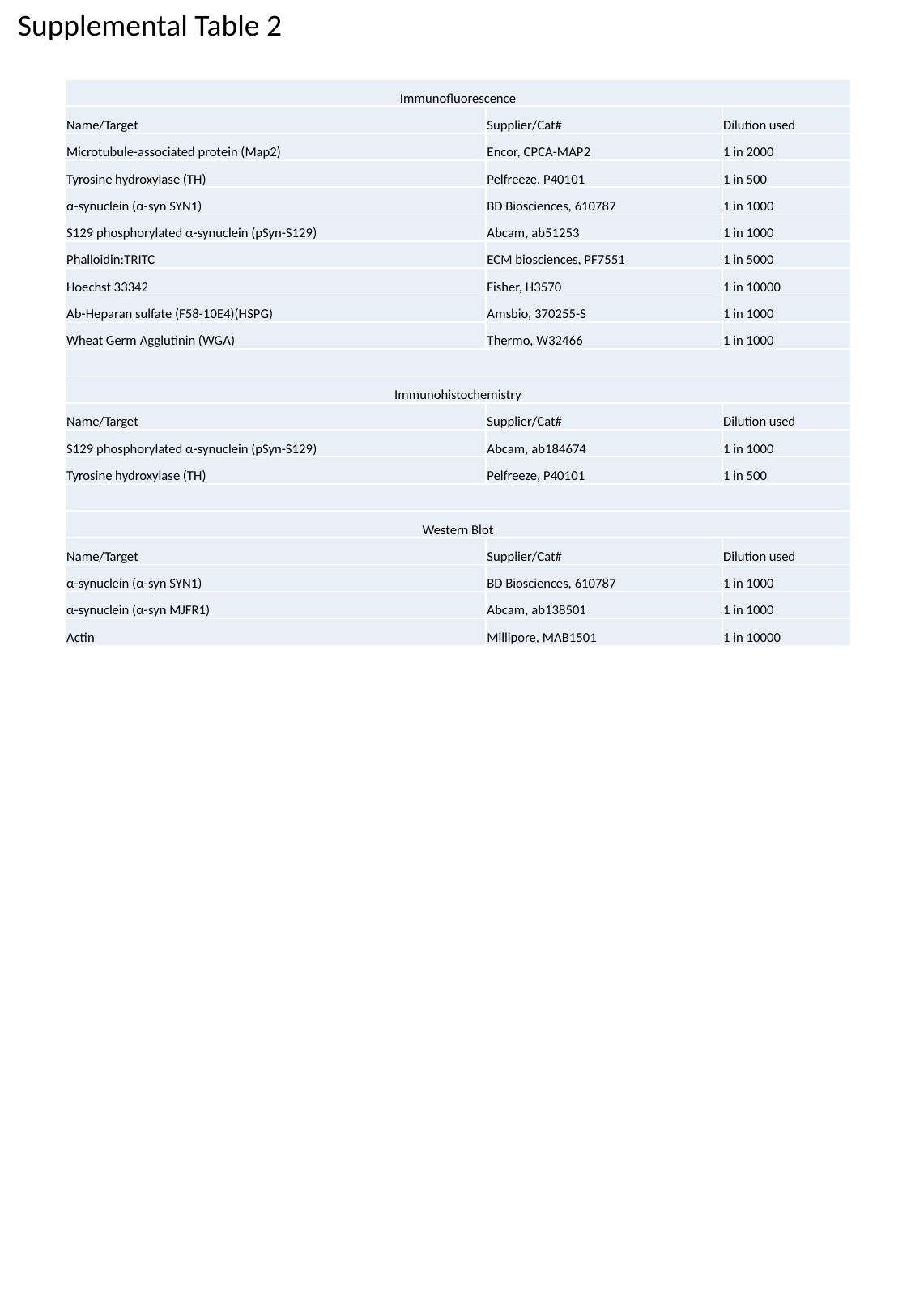

Supplemental Table 2
| Immunofluorescence | | |
| --- | --- | --- |
| Name/Target | Supplier/Cat# | Dilution used |
| Microtubule-associated protein (Map2) | Encor, CPCA-MAP2 | 1 in 2000 |
| Tyrosine hydroxylase (TH) | Pelfreeze, P40101 | 1 in 500 |
| α-synuclein (α-syn SYN1) | BD Biosciences, 610787 | 1 in 1000 |
| S129 phosphorylated α-synuclein (pSyn-S129) | Abcam, ab51253 | 1 in 1000 |
| Phalloidin:TRITC | ECM biosciences, PF7551 | 1 in 5000 |
| Hoechst 33342 | Fisher, H3570 | 1 in 10000 |
| Ab-Heparan sulfate (F58-10E4)(HSPG) | Amsbio, 370255-S | 1 in 1000 |
| Wheat Germ Agglutinin (WGA) | Thermo, W32466 | 1 in 1000 |
| Immunohistochemistry | | |
| Name/Target | Supplier/Cat# | Dilution used |
| S129 phosphorylated α-synuclein (pSyn-S129) | Abcam, ab184674 | 1 in 1000 |
| Tyrosine hydroxylase (TH) | Pelfreeze, P40101 | 1 in 500 |
| Western Blot | | |
| Name/Target | Supplier/Cat# | Dilution used |
| α-synuclein (α-syn SYN1) | BD Biosciences, 610787 | 1 in 1000 |
| α-synuclein (α-syn MJFR1) | Abcam, ab138501 | 1 in 1000 |
| Actin | Millipore, MAB1501 | 1 in 10000 |
